# A Gamma-adapted recombinant subunit vaccine induces broadly neutralizing antibodies against SARS-CoV-2 variants and protects mice from infection

**DOI:** 10.1101/2023.01.03.522213

**Authors:** Lorena M. Coria, Juan Manuel Rodriguez, Agostina Demaria, Laura A. Bruno, Mayra Rios Medrano, Celeste Pueblas Castro, Eliana F. Castro, Sabrina A. Del Priore, Andres C. Hernando Insua, Ingrid G. Kaufmann, Lucas M. Saposnik, William B. Stone, Lineia Prado, Valeria Krum, Francisco M. Zurvarra, Johanna E. Sidabra, Ignacio Drehe, Jonathan A. Baqué, Mariana Li Causi, Analia V. De Nichilo, Cristian J. Payes, Albert J. Auguste, Julio C. Vega, Diego E. Álvarez, Juan M. Flo, Karina A. Pasquevich, Juliana Cassataro

## Abstract

The COVID-19 pandemic continues with the emergence of successive new variants of concern (VOC). One strategy to prevent breakthrough infections is developing safe and effective broad-spectrum vaccines. Here, we present preclinical studies of a RBD recombinant vaccine candidate derived from the Gamma SARS-CoV-2 variant adjuvanted with alum. Gamma RBD-derived antigen elicited better neutralizing antibody and T cell responses than formulation containing ancestral RBD antigen. The Gamma-adapted subunit vaccine elicited a long-lasting antibody response with cross-neutralizing activity against different VOC including the Omicron variant. Additionally, Gamma variant RBD-adapted vaccine elicited robust T cells responses with induction of Th1 and CD8^+^ T cell responses in spleen and lung. Vaccine-induced immunity protected K18-hACE2 mice from intranasal challenge with SARS-CoV-2 increasing survival, reducing body weight loss and viral burden in the lungs and brain. Importantly, the subunit vaccine demonstrated a potent effect as heterologous booster of different vaccine platforms including the non-replicating adenovirus vaccine ChAdOx1-S, the mRNA vaccine BNT162b2 and the inactivated SARS-CoV-2 vaccine BBIBP-CorV, increasing cross-reactive antibody responses. Our study indicates that the adjuvanted Gamma RBD vaccine is highly immunogenic and a broad-spectrum vaccine candidate to combat SARS-CoV-2 variants including Omicron.

## Introduction

Severe Acute Respiratory Syndrome Coronavirus 2 (SARS-CoV-2) has infected over 630 million people worldwide and resulted in more than 6,6 million deaths (WHO. World Health Organization, Weekly Operational Update on COVID-19). The vaccines authorized for emergency use or fully approved are safe and highly effective against severe disease^1–5^. However, it has been reported that vaccine-induced protection against symptomatic SARS-CoV-2 infection wanes over time^6,7^. Additionally, as COVID-19 pandemic progresses several variants of concern (VOCs) have emerged, including B.1.351 (Beta), B.1.617.2 (Delta) and P.1 (Gamma), B.1.1.529 (Omicron BA.1) and its subvariants (BA.2.12.1, BA.4/BA.5, BQ.1 and XBB and its descendants). Omicron lineages have been reported to be more resistant to neutralization by vaccine-elicited antibodies and, in some cases are more transmissible than previous VOCs^8–12^

Wanning of vaccine-induced immunity and antibody evasive virus variants create the need for new vaccine strategies that can induce more potent, durable, and broader immune responses to enhance protection against SARS-CoV-2. Additional booster doses of current vaccines have been implemented around the world. Current approved vaccines are mainly directed against the spike protein of the prototype Wuhan-1 SARS-CoV-2 strain therefore their efficacy against certain VOCs is limted^11^. Therefore, vaccines are being adapted to variants as a strategy to improve vaccine effectiveness. While current vaccines may be effective at reducing severe disease to existing VOCs, variant-specific antigens, whether in a mono- or multivalent-vaccine, may be required to induce optimal immune responses and reduce infection against emerging variants.

Most of the variant-adapted vaccines boosters were bivalent or monovalent containing beta and ancestral spike antigens besides Omicron^13–16^. Interestingly, the Beta variant originated in South Africa, while the Gamma variant originated in Brazil a few months later. Until now, there are few studies using a Gamma variant adapted vaccine ^17–19^.

In this work, we describe the design, formulation, and immunogenicity of a RBD vaccine candidate comparing ancestral, and Gamma derived RBD versions. Our results show that Gamma-adapted RBD vaccine is more immunogenic than ancestral RBD vaccine in terms of inducing broader neutralizing antibodies (and higher antigen (Ag) specific cellular immune responses. Therefore, we selected RBD Gamma as antigen to develop a recombinant subunit vaccine that can be used as primary or booster vaccine. Here we describe the rational design, generation, and preclinical evaluation of the Gamma RBD-based vaccine candidate using aluminum hydroxide as adjuvant. Immunogenicity was evaluated after a primary two-dose vaccine schedule and after a heterologous booster of different anti-SARS-CoV-2 vaccine platforms including the non-replicating adenovirus vaccine ChAdOx1-S (Oxford AstraZeneca), the mRNA vaccine BNT162b2 (Pfizer) and the inactivated SARS-CoV-2 vaccine BBIBP-CorV (Sinopharm).

## Results

### Design and production of RBD antigens

To compare immune responses elicited with ancestral or Gamma derived RBD, two formulations were developed using RBD homodimers from the ancestral (Wuhan-Hu-1) and Gamma variants. Antigens comprise single-chain dimeric repeats of the RBD protein from amino acids 319R to 537K. The Gamma RBD antigen includes the mutations described in the Gamma SARS-CoV-2 variant of spike in this region: K417T; E484K and N501Y (P.1/501Y.V3). High-productivity clones were generated under the genetic background of a CHO-S (Chinese Hamster Ovary) cell line in high-density suspension cell cultures. Then, proteins were purified, and antigen purity confirmed by SDS-PAGE and Western Blot (**Fig. 1A–B**). Comparison of the binding affinity of Gamma and ancestral RBD to hACE2 receptor was performed by ligand-receptor binding ELISA (**Fig. 1C**). Both antigens bind to hACE2 at comparable levels with a slight greater binding affinity of Gamma RBD. Purified antigens were then absorbed to aluminum hydroxide (alum) to generate the final formulations.

**Figure 1.**
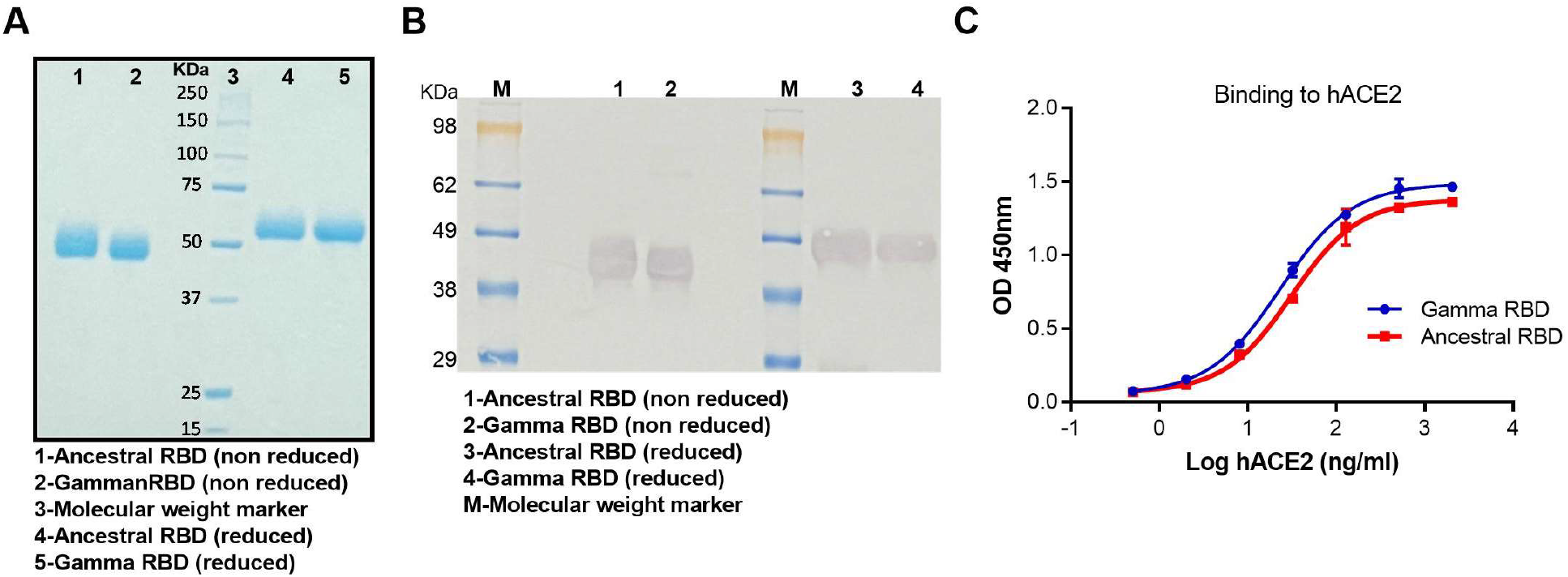
Characterization of RBD antigens. **A**. Non-reduced (nr) and reduced (r) SDS-PAGE migration profiles of the pooled samples (Gamma and ancestral RBD) are shown. **B**. **Western blot.** Non-reduced and reduced RBD antigens were detected using a rabbit polyclonal anti-RBD antibody. **C**. Binding affinity of Gamma and ancestral RBD proteins to immobilized human ACE2 receptor by ligand-receptor binding ELISA. Experiments were performed in duplicates and mean ± SD values are shown.

### Gamma RBD antigen vaccine formulation improved antibody and cellular immune responses in comparison with ancestral RBD antigen

Immunogenicity of vaccine formulations was assessed in BALB/c mice after i.m. immunization in a two-dose vaccine schedule using 10μg of each antigen absorbed in aluminum hydroxide (**Fig. 2A**). RBD (Gamma and ancestral)-specific IgG was evaluated by ELISA in sera of immunized animals. Both formulations induced high levels of IgG antibodies against the ancestral and Gamma RBD antigens (**Fig. 2B**). Anti-RBD IgG responses elicited by the formulation containing the Gamma antigen were significatively higher than that induced by the ancestral RBD-based vaccine on days 14 and 42 after prime immunization (**Fig. 2C**).

**Figure 2.**
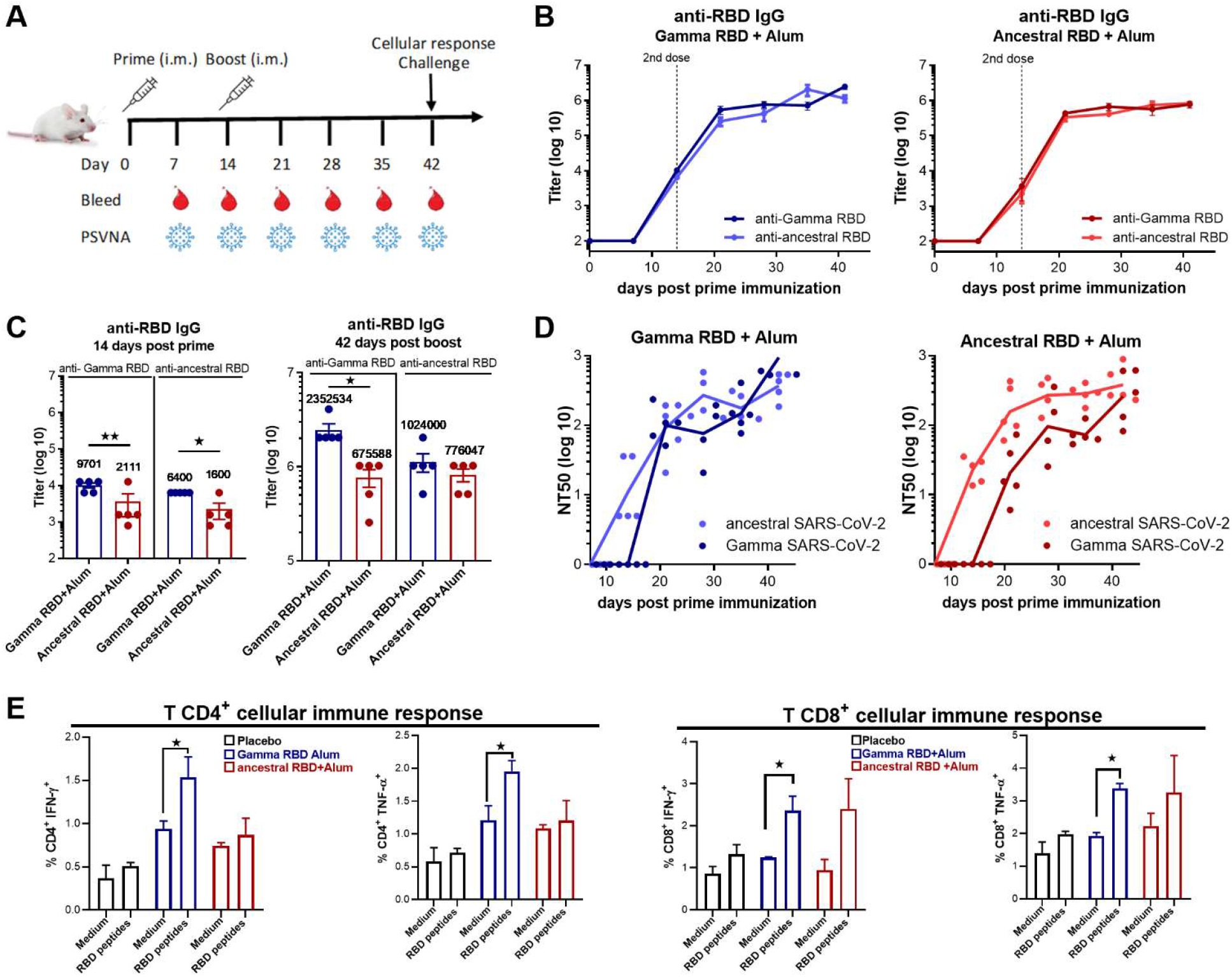
Gamma RBD-adapted vaccine induce antibodies with a broad neutralizing activity and Ag-specific T cell responses. **A**. Immunization protocol scheme. BALB/c mice were immunized at day 0 and day 14 via i.m. with: ancestral RBD + Alum (n=5), Gamma RBD + Alum (n=5). Serum samples were obtained at indicated time points for ELISA and neutralization assays. **B**. Kinetics of RBD-specific IgG endpoint titers in sera of immunized animals by ELISA. Points are means ± SEM. **C**. RBD-specific IgG titers in sera of immunized animals at day 14 and 42 post prime immunization. *p<0.05. **p<0.01. Mann Whitney test. **D**. Neutralizing-antibody titers in sera was determined by authentic PRNT SARS-CoV-2 assay for each group of vaccinated mice at different time points. The black lines represent the geometric mean (GMT) of all data points. Titers correspond to the 50% of virus neutralization (NT50). **E**. Mice were sacrificed 42 days after the first immunization to obtain spleens and lungs and T cell response was evaluated by intracellular flow cytometry. Splenocytes were stimulated with complete medium or RBD-peptides pool for 18 h and then brefeldin A was added for 5 h. Afterward, cells were harvested and stained with specific Abs anti-CD8, and anti-CD4, fixed, permeabilized, and stained intracellularly with anti–IFN–γ and anti-TNF-α. Results are presented as percentage of IFN-γ or TNF-α – producing T lymphocytes. Bars are means ± SEM. *p < 0.05 vs. medium. T test.

Neutralizing antibody activity was evaluated by a live virus assay using ancestral or Gamma SARS-CoV-2 variants. Immunization with the formulation containing ancestral RBD antigen induced higher neutralization titers against ancestral SARS-CoV-2 than against Gamma variant (**Fig. 2D**, p<0.05 NT50 Gamma vs ancestral, time points from 14 to 35 days post prime immunization). In contrast, vaccination with Gamma RBD plus Alum elicited high neutralizing antibody (Ab) titers against both viruses. In addition, cellular immune responses were analyzed in splenocytes from immunized animals by intracellular flow cytometry. Gamma RBD-based vaccine induced higher frequency of IFN-y and TNF-α producing CD4^+^ and CD8^+^ T cells (p<0.05, medium alone vs RBD peptides stimulation) compared to the frequency induced by the formulation with the ancestral RBD antigen (**Fig. 2E**). These results demonstrated that vaccine containing Gamma RBD antigen induces a broader neutralizing response and a superior Ag-specific T cellular immune response than RBD derived from the ancestral SARS-CoV-2. Thus, we decided to move forward with the formulation containing the Gamma RBD antigen.

### Gamma RBD adjuvanted formulation induces long-lasting antibody responses with broad neutralizing activity against different virus variants of concern

Binding antibody responses were evaluated over longer periods after mice immunization revealing that anti-RBD IgG titers remained high up to 253 days (more than 8 months) post prime immunization (**Fig. 3B**). Additionally, neutralization capacity of antibodies elicited after immunization was evaluated using different VOCs in a live virus PRNT assay at this time point. Immunization with Gamma RBD adsorbed to Alum elicited high neutralizing Ab titers against Gamma (GMT 254.8) and ancestral (GMT 312.8) virus. Interestingly, elicited antibodies showed cross-reactivity against Alpha (GMT 67.82), Delta (GMT 85.49), Omicron BA.1 (GMT 135.5) and BA.5 (GMT 33.12) VOCs (**Fig. 3B**). Of note, there were no significant differences between these variants indicating that Gamma RBD vaccination elicits broadly long-lasting neutralizing Ab responses.

**Figure 3.**
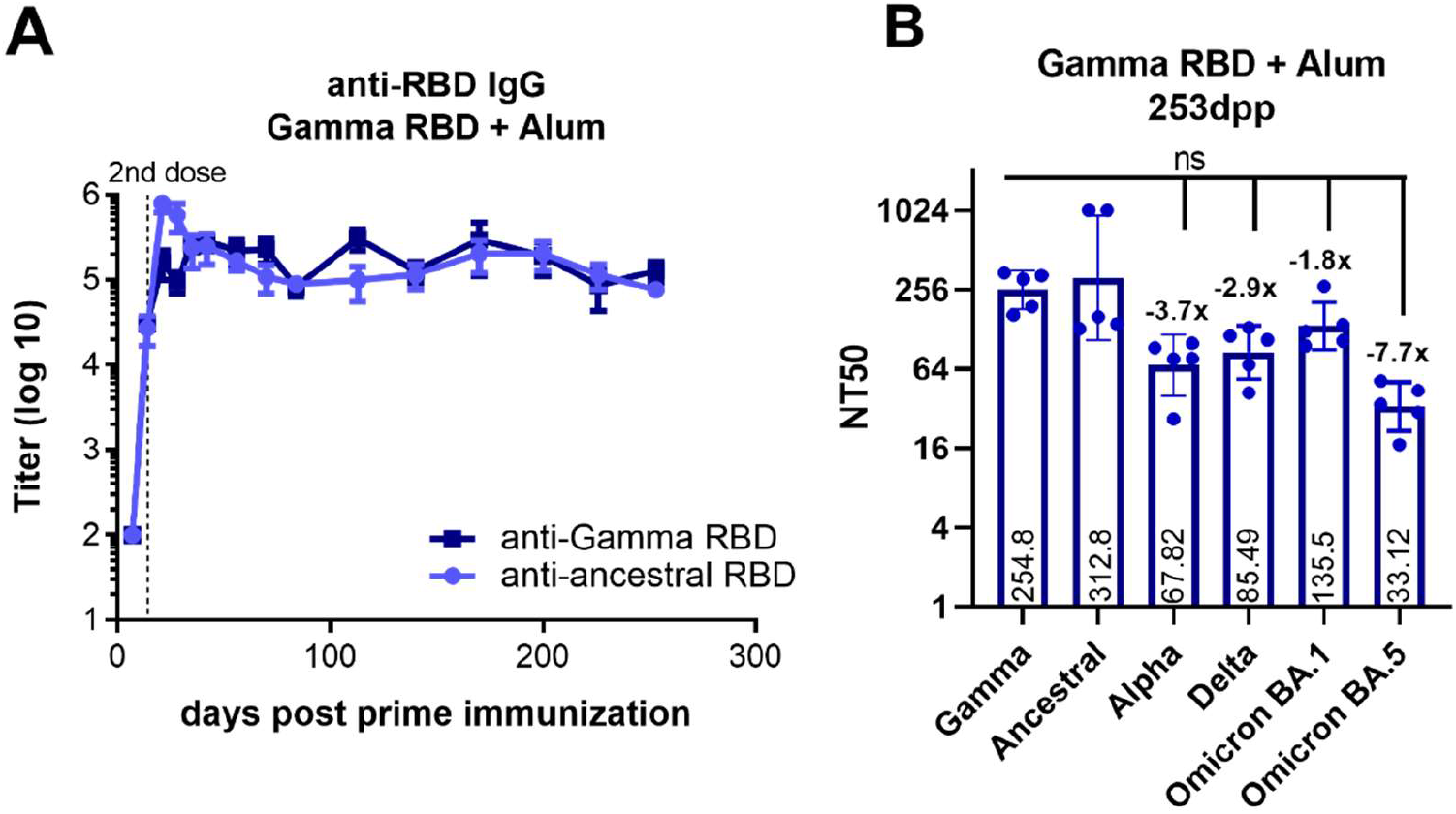
Gamma RBD plus alum vaccinated mice induce a long-lasting antibody response with broaden neutralization activity. BALB/c mice were immunized at day 0 and day 14 via i.m. with: Gamma RBD + Alum (n=5). **A**. Kinetics of RBD-specific (ancestral or Gamma) IgG endpoint titers in sera of immunized animals by ELISA until 253 days post prime immunization (dpp). Points are means ± SEM. **B.** Neutralizing-antibody titers against ancestral SARS-CoV-2 and different VOC (Alpha (B.1.1.7), Gamma (P.1) Delta (B.1.617.2) and Omicron (BA.1 and BA.5) were evaluated 253dpp in Gamma RBD +Alum immunized group. Neutralization titer was defined as the serum dilution that reduces 50% the cytopathic effect (NT50).

### Gamma RBD-adapted vaccine elicits Ag-specific mixed Th1-Th2 immune responses

Figure 2 shows that Gamma RBD adjuvanted formulation induces Ag-specific T cell responses in splenocytes. The cell-mediated immune responses in spleen and lung generated after vaccination was further evaluated. Gamma RBD plus alum vaccination induced Ag-specific IFN-y and IL-5 secretion in spleen and lung cells (**Fig. 4A**) indicating that a mixed Th1/Th2 is generated at local and systemic level.

**Figure 4.**
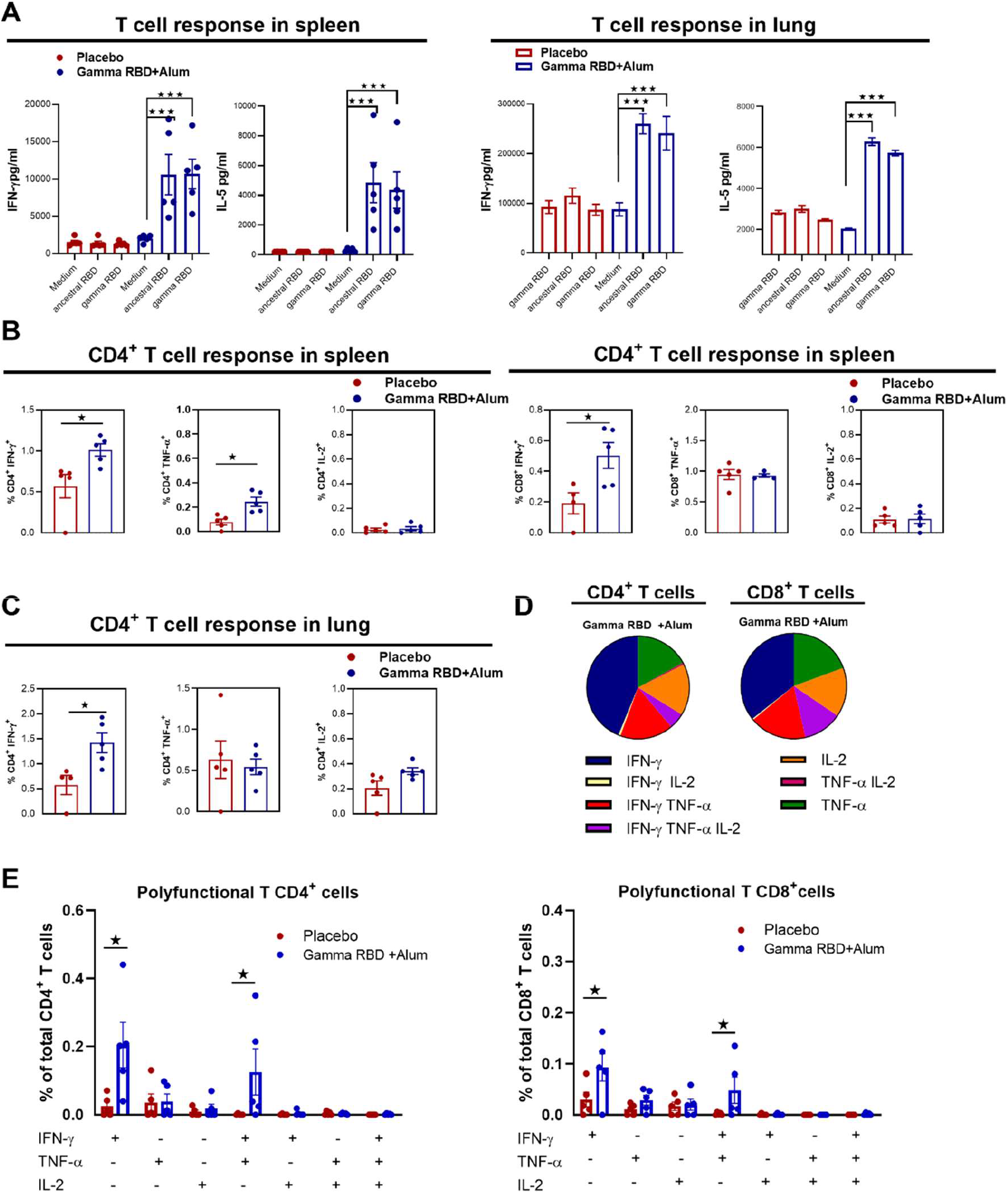
Gamma RBD formulated with alum induces T cell responses in mice. Cellular response was evaluated in spleen and lung 28 days after last immunization. **A**. Frequency of secreted IFN-y and IL-5 following splenocytes or lung cells stimulation with medium or recombinant RBD were determined by ELISA. Bars are means ± SEM of pg/ml of IFN-y and IL-5 after subtracting the amount in medium stimulated cells. *p < 0.05, **p < 0.01. T test. **Intracellular flow cytometry analysis of cytokine secreting T cells**. Splenocytes (**B**) and lung cells (**C**) were stimulated with complete medium or a peptide pool derived from RBD and then brefeldin A was added. Afterward, cells were harvested and stained with specific Abs anti-CD8, and anti-CD4, fixed, permeabilized, and stained intracellularly with anti-IFN-γ, TNF-α and anti-IL-2. Results are presented as percentage of cytokine-producing T cells. Bars are means ± SEM. *p < 0.05 vs. medium. T test. **D**. **Assessment of polyfunctional T cells**. Frequency of CD4^+^ and CD8^+^ T cells population from splenocytes that produce one, two or three cytokines were plotted for each group of mice. **E**. Pie graphs indicate the total proportion of RBD specific cytokine production in splenocytes from the Gamma RBD + Alum immunized mice.

In the spleen, antigen-specific IFN-y- and TNF-α-producing CD4^+^ T cells and IFN-y producing CD8^+^ T cells were detected upon Ag stimulation of cells from vaccine immunized group (**Fig. 4B**). In the lung, immunization with Gamma RBD + Alum elicited RBD-specific IFN-y producing CD4^+^ T cells (**Fig. 4C**). Production of cytokines by CD8^+^ T cells was not detected in the lungs (data not shown).

A large proportion of antigen-specific cytokine producing CD4^+^ T and CD8^+^ T cells in spleens were double positive IFN-y^+^ TNF-α^+^ and single positive IFN-y^+^ Th1 cells (**Fig. 4D–E**). Analysis of total CD4^+^ and CD8^+^ T cell population that produce cytokines revealed that induced multifunctionality is greater in the population of CD8^+^ T cells (**Fig. 4E**).

These results suggest that the Gamma RBD-based vaccine promotes a specific T cell response biased to a multifunctional Th1 profile and effector CD8^+^ T cells able to produce IFN-y in spleen. Interestingly, a Th1 response with production of IFN-y was observed in the lungs.

### Gamma RBD-based vaccine protects mice after intranasal SARS-CoV-2 challenge

To determine vaccine ability to protect against SARS-CoV-2 infection, a lethal K-18-hACE2 transgenic mice model was used^20,21^. Immunized mice were challenged intranasally with 2×10^5^ PFU of SARS-CoV-2 and monitored daily for body weight changes and survival, a subset of mice was euthanized on day 5 post-infection (p.i.) to measure viral loads in the lungs and brains.

Body weight started to decrease in infected placebo mice after 3 days p.i. reaching a 20% body weight decrease on day 9 p.i. whereas mice immunized with the vaccine did not show significant weight loss during the experiment (**Fig. 5A**). Moreover, 80% of infected placebo mice died approximately 6 days p.i., 83% of vaccine immunized mice where alive at this time point. (**Fig. 5B**). After 9 days, all mice in the placebo group succumb to infection whereas a 66% of the immunized mice survived (P = 0.0208 vaccinated vs placebo).

**Figure 5.**
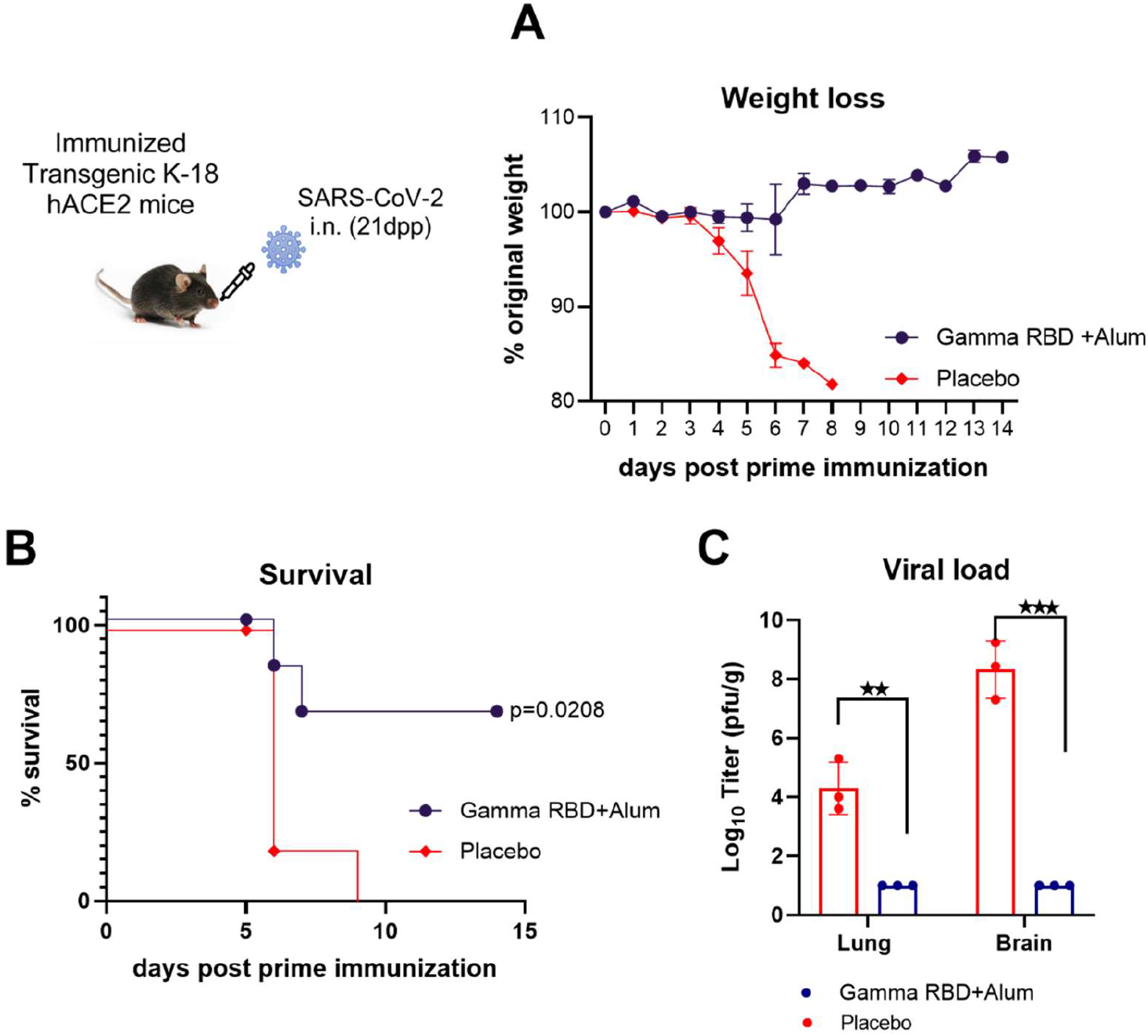
Vaccination with Gamma RBD + Alum protects K18-hACE2 transgenic mice against intranasal SARS-CoV-2 infection. Mice received PBS (Control) (n=8) or Gamma RBD + Alum (n=9) administered via i.m. route at day 0 and 14. Four weeks following immunization, K18-hACE2 mice were intranasally infected with 2 × 10^5^ PFU of SARS-CoV-2. **A**. Weight changes in mice were monitored daily until day 14 after infection. Points are means ± SEM of percentage of original weight. **B**. Mice survival were monitored daily until day 14. Each dot represents the percentage of mice alive at that time point. Survival curves were analyzed with Log-rank (Mantel-Cox) test. P=0.0208 vs placebo. **C**. Five days after infection lungs and brains (n=3) were obtained from groups of mice and SARS-CoV-2 virus was titrated. Bars represent the mean ± SEM. **p < 0.01 and ***p < 0.001. T test.

Five days post infection, a subset of animals was euthanized to assess viral loads in the lungs and brains. High infectious SARS-CoV-2 titers were observed in the lungs and brains of the placebo group of mice while low virus titers were detected in brains of animals vaccinated with Gamma RBD + Alum (**Fig. 5C**). Thus, the Gamma RBD-based vaccine induced protection against experimental intranasal challenge with SARS-CoV-2 in a mouse model of severe disease.

### A booster dose of Gamma RBD-based vaccine induces higher neutralizing Ab titers than homologous booster vaccination in mice previously primed with different vaccine platforms

Booster vaccination has become a strategy to prevent SARS-CoV-2 breakthrough infections. Immunogenicity of the Gamma RBD adjuvanted vaccine as a heterologous booster for different primary vaccine platforms was studied in comparison with homologous regimens (i.e., three dose of the same vaccine). To this end, each group of mice was immunized with one of the following vaccine platforms in a two-dose scheme primary vaccination: i) non-replicating adenovirus vaccine ChAdOx1-S (Oxford AstraZeneca), ii) the mRNA vaccines BNT162b2 (Pfizer) and iii) the inactivated SARS-CoV-2 vaccine BBIBP-CorV (Sinopharm). Three months later, half of the animals received a homologous boost and the other half a heterologous boost with the Gamma RBD-based vaccine (**Fig. 6A**).

**Figure 6.**
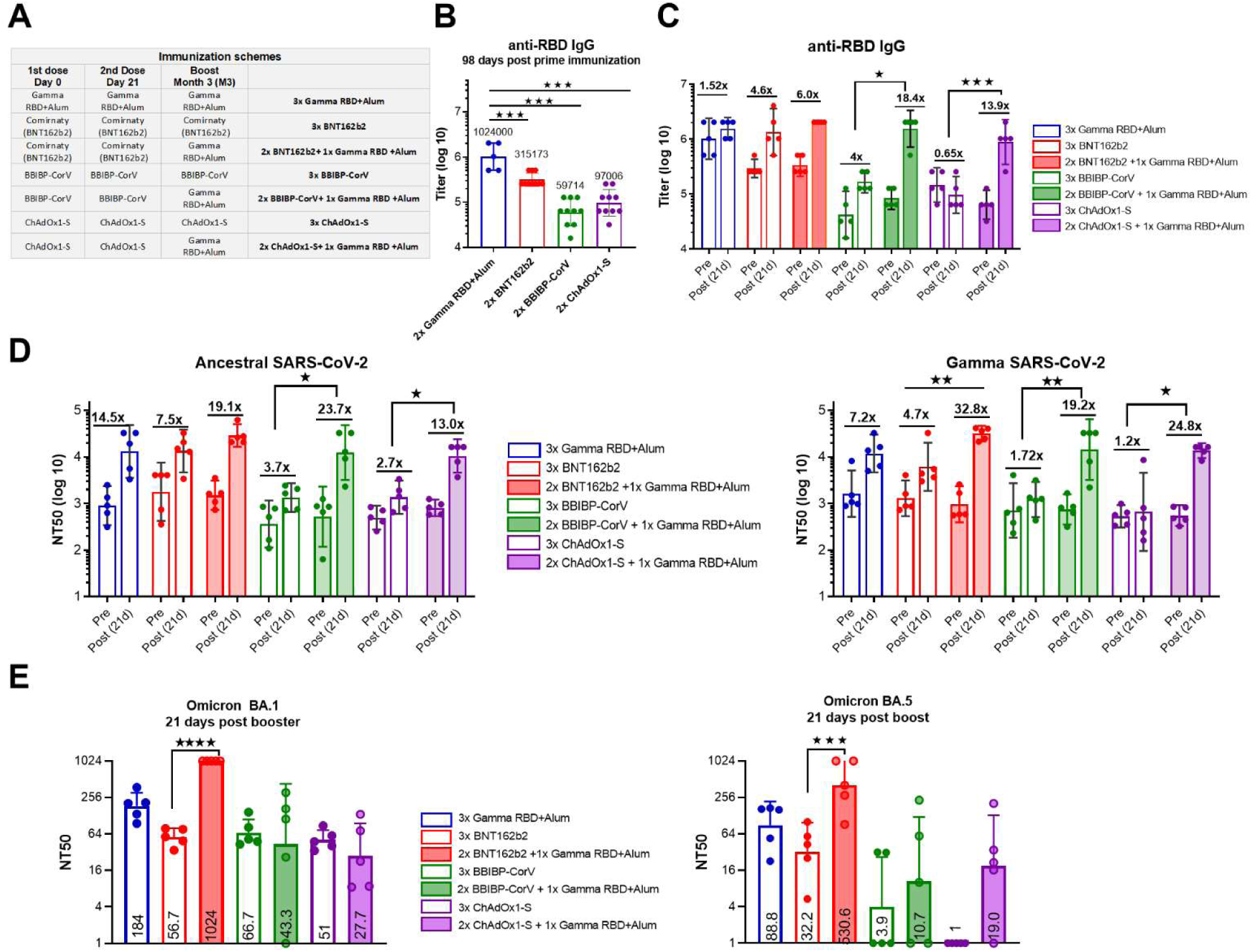
Heterologous boosting with Gamma RBD-adapted vaccine formulation of different primary vaccine platforms. **A**. Immunization protocol scheme. BALB/c mice were immunized at day 0 and day 21 via i.m. with: Gamma RBD + Alum (n=5), ChAdOx1-S (n=10), BNT162b2 (n=10) and BBIBP-CorV (n=10). At day 100 (M3) half of mice received homologous third doses (BNT162b2, BBIBP-CorV or ChAdOx1-S accordingly to table in A) and the other half heterologous booster shots with Gamma RBD + Alum. **B**. RBD-specific IgG titers in sera of immunized animals at day 98 post prime immunization. Bars are GMT ± 95% CI. *p<0.05. **p<0.01. Kruskal Wallis. Bars are GMT ± 95% CI. Numbers in bold are fold-increase in GMT values. **C**. Comparison of RBD-specific IgG titers pre and post-boost immunization. Numbers in bold are GMT fold-increase values. Bars are GMT ± 95% CI *p<0.05. ***p<0.001. Kruskal Wallis. **D**. Neutralizing-antibody titers against ancestral and Gamma SARS-CoV-2 were evaluated pre (98dpp) and post boost immunization (121 dpp) of mice. Neutralization titer was defined as the serum dilution that reduces 50% the cytopathic effect (NT50). Bars are GMT ± SD. Numbers in bold are fold-increase GMT values for each group. *p<0.05. ***p<0.001. Kruskal Wallis test. **E**. Neutralizing activity of antibodies against omicron (BA.1 and BA.5) were evaluated 21 days post boost immunization of mice. Neutralization titer was defined as the serum dilution that reduces 50% the cytopathic effect (NT50). Bars are GMT ± SD and numbers are GMT values for each group of mice. *p<0.05. ***p<0.001. Kruskal Wallis test.

All vaccine platforms induced RBD-specific IgG after two dose primary vaccination (**Fig 6B**). Interestingly, three months after primary vaccination the Gamma RBD-based vaccine candidate elicited the highest Ab titers (day 98 post prime immunization, **Fig. 6B**). Three weeks after the third dose, in the group that received a primary regimen with the inactivated BBIBP-CorV-vaccine the increase in RBD-specific IgG titers was higher after homologous boosting with Gamma-adapted vaccine than with the homologous vaccine (18.4× vs. 4×-fold-increase respectively, p<0.05). Similarly, primary vaccination with the non-replicating adenovirus vaccine group ChAdOx1-S and booster with Gamma RBD vaccination showed a higher fold-increase in anti-RBD IgG titers than homologous ChAdOx1-S boosting (13.9× vs. 0.65×-fold-increase respectively, p<0.001. **Fig. 6C**). In the group that received mRNA vaccine as primary regimen the increment in RBD-specific IgG titers was similar after homologous or heterologous booster (6× vs. 4× fold rise respectively, p>0.05. **Fig. 6C**).

Analysis of the neutralizing activity of sera from mice before (pre) and after booster (post) immunization against ancestral SARS-CoV-2 revealed that the fold-increase in neutralizing Ab titer were significantly higher after heterologous booster with the Gamma subunit vaccine than homologous booster with BBIBP-CorV and ChAdOx1-S vaccines (**Fig. 6D,** p<0.05). Heterologous boosting with the Gamma subunit vaccine induced a significantly higher increase of the neutralizing Abs titers against the Gamma VOC than homologous boosting in all vaccine platforms used to prime mice (**Fig. 6D**). The best neutralization activity (against ancestral and Gamma variants) was observed in the group that received primary immunization with the mRNA vaccine BNT162b2 and boosted with the Gamma RBD-adapted vaccine (**Fig 6D**).

Finally, cross-neutralization activity was assessed against Omicron subvariants BA.1 and BA.5 three weeks after the booster dose. In all groups, antibody neutralization capacity was reduced against Omicron subvariants compared to neutralizing Ab titers against the ancestral and Gamma variant. However, a degree of cross-reactive antibodies against BA.1 and BA.5 was detected, and the groups boosted with the subunit vaccine outperformed groups boosted with homologous vaccines (adenoviral, inactivated and mRNA vaccines, **Fig. 6 D–E**). In particular, in the group that received mRNA as primary vaccination, the neutralizing Ab titers against Omicron BA.1 and BA.5 were significantly higher after a heterologous boost with Gamma RBD-adapted subunit vaccine than with a homologous boost vaccination **(Fig. 6D–E)**.

These results suggest that the Gamma RBD-adapted recombinant vaccine candidate could serve as an effective booster of different primary vaccine platforms inducing a broad cross-reactive immunity.

## Discussion

The rapid emergence of SARS-CoV-2 variants has challenged the global development of vaccines. The ability of VOCs to escape neutralization by antibodies elicited by infection, vaccination, or therapeutic application has been widely studied^22,23^. Developing variant adapted vaccines that can elicit broad and strong immune responses against these predominant circulating variant(s) is proposed as one of the best strategies to address variants immune evasion.

In this study, we described the immunogenicity of a recombinant adapted subunit vaccine formulated with aluminum hydroxide to be used as a primary or booster vaccine. Ancestral and Gamma variant derived RBD antigens were compared in their capacity to induce neutralizing Ab responses against SARS-CoV-2. It was reported that sera from individuals vaccinated with an ancestral RBD antigen plus alum vaccine formulation showed a reduction in neutralizing antibody titers of 1.7x and 2.1x fold against Gamma and Beta variants of SARS-CoV-2 respectively^24^. In agreement with these results ancestral RBD could not induce good neutralizing activity against the Gamma SARS-CoV-2 variant whereas we demonstrated for the first time that the vaccine formulation containing Gamma RBD was able to induce neutralizing Abs against ancestral and Gamma SARS-CoV-2 variants.

Beta and Gamma variants, which originated in South Africa and Brazil, respectively, carry the K417N/T, E484K and N501Y amino acid changes in the spike RBD. Mutations in the Beta variant decreased the neutralization sensitivity of convalescent and vaccinated sera by 6x-fold, while mutations in Gamma decreased neutralizing activity by 3x-fold ^25^. Mono and bivalent candidates containing different combinations of VOCs based antigens have been assessed in humans ^13,26^. Omicron-adapted mRNA vaccines as boosters have been implemented in several countries. Recently, a Beta-adapted subunit vaccine has been approved for use in Europe. Until present, there are no clinical studies with a Gamma adapted vaccine.

The Gamma variant booster vaccine candidate tested here showed a broad cross-reactive neutralizing antibody response against several VOCs including Omicron in mice. This could be related to the notion that K417N/T and E484K substitutions shared by Beta and Gamma viruses simulate a structure that enhances the spectrum of neutralizing activity. In agreement with this hypothesis and our results, a study evaluating the immunogenicity of an S-protein sequence containing these three substitutions in addition to D614G (B.1) demonstrated that the variant vaccine is capable of eliciting high neutralizing Ab titers that cross-reacted with five VOCs in NHP^27^. Moreover, in preclinical and clinical trials Beta derived vaccines induced broadly neutralizing antibodies that can cross-react against different VOC ^27,28^ in a similar way than the Gamma derived vaccine presented in this work.

A recombinant vaccine containing RBD-dimers derived from ancestral SARS-CoV-2 and aluminum hydroxide as adjuvant proved to be safe and immunogenic in humans^29^. Structural analysis of these RBD-dimers indicates that are arranged in a way where the receptor binding motif (RBM) is correctly presented and the major antigenic sites of RBD are exposed^30^. This supports the versatility of the RBD-dimers as a module to adapt different variants for the induction of broader immune responses. Hence, it has been reported that chimeric RBD-dimers elicit broader responses to variants compared with homodimers^30^. Increasing evidence suggests that vaccine protection against COVID-19 might be waning over time, and newly emerging variants, such as Delta and Omicron, might evade vaccine-induced immune protection. Our vaccine induced a long-lasting antibody response with neutralizing activity against different VOCs up to 253 days after the last immunization. Adjuvanted recombinant COVID-19 vaccine formulations have reported durability of the neutralizing-antibody responses for 7 months post-immunization in NHP^31^. This study together with previous reports suggest that recombinant variant-specific vaccines may be needed for improved and long-lasting protection against known and emerging VOCs.

Main correlates of protection against COVID-19 are binding and neutralizing antibody titers^32^. However, cellular immunity is associated with positive outcomes^33^. Here, we have provided a comprehensive assessment of the T cell response to a Gamma-adapted recombinant subunit vaccine. Gamma RBD + alum vaccination elicited a robust Th1-dominated T cell response and RBD-specific CD8^+^ T cell response. Analysis of polyfunctionality on vaccine elicited T cells suggests that the major proportion of cells produce one or two cytokines (IFN-γ and TNF-α). Polyfunctional Th1 responses were reported after human vaccination with adenoviral ChAdOx1-S and the mRNA BNT162b2 vaccine^34,35^. Protection studies in transgenic mice presented here confirm that the immunity induce by the Gamma-derived RBD vaccine candidate can protect mice from a lethal nasal challenge of ancestral SARS-CoV-2 virus.

Updated immunization strategies were implemented as booster shots across the globe. In this work, we showed that vaccination with Gamma RBD plus alum as booster of adenoviral, inactivated or mRNA-based vaccines induced a higher binding and neutralizing antibody titer increase against different VOCs than vaccinations with homologous vaccines. In general, heterologous boosters showed higher vaccine effectiveness than homologous boosters for all outcomes in humans, providing additional support for a mix-and-match approach^36,37^. Our vaccine not only increased GMT values of neutralizing Ab against ancestral and Gamma virus variants but also against Omicron lineages (BA.1 and BA.5).

The Gamma-RBD vaccine candidate reported in this article has demonstrated to be safe in non-clinical safety studies. Both the antigen and the formulated vaccine are being produced under GMP at manufacturing plants of Laboratorio Pablo Cassará S.R.L. located in Buenos Aires. Stability studies showed that the vaccine can be stored and handled in refrigerated conditions (2 – 8 °C) and does not need freezing temperatures. The development of a vaccine does not require stringent cold-chain transport and storage facilitates its availability at local and global supply. This gamma RBD plus alum vaccine was named ARVAC Cecilia Grierson and is currently in a Phase 1 clinical trial (NCT05656508).

This work highlighted the value of a Gamma variant-adapted vaccine to induce higher and broader immune responses against SARS-CoV-2 at booster vaccination and set the precedent for its further clinical development.

## Methods

### Antigen expression and purification

Constructs from Wuhan (ancestral) and Gamma SARS-CoV-2 variants of the antigen consisted of a single-chain dimer of the receptor binding domain (RBD), comprising amino acids 319R to 537K of the Spike protein from ancestral and Gamma virus variant. The signal peptide sequence from the SARS-CoV-2 Spike protein was used for protein secretion. The constructs were codon-optimized for CHO cell expression and synthesized by GenScript (Hong Kong Limited, Japan). All constructs were cloned into the pJC3UMCS-4 expression vector (Pablo Cassara Foundation) comprising a CMV promoter and a cis-acting sequence for minimized host gene silencing. Linearized forms of the vectors were transfected into CHO-S cells by lipofection. After 15-20 days of G418 antibiotic selection, an end point dilution procedure was applied for clone isolation. Clone candidates were selected by specific and volumetric productivity assessed by ELISA. Recombinant protein relative abundance was confirmed by SDS-PAGE and Coomasie Brilliant Blue G-250 staining.

Pilot batches of antigen and vaccines were manufactured by Laboratorio Pablo Cassará, Argentina. Controlled cell substrates were used to inoculate a 2-L bioreactor with 1.4 L as working volume. An alternate tangential flow system (ATF, Repligen) was used for perfusion. The harvest after 20 - 25 days was used as starting material for a downstream process based on three chromatographic steps: first an affinity mix-mode capture chromatography step followed by two different ionic chromatography to eliminate residual host cell DNA and host cell proteins and other process-related impurities. TFF (Tangential flow filtration) was used to adjust pH, conductivity, and protein concentrations of the batches.

### SDS-PAGE and western blot analysis

RBD samples were run under reducing and non-reducing conditions by SDS-PAGE (10% polyacrylamide). The samples were visualized by staining with Coomassie Brilliant Blue G-250. Bands were then transferred to a nitrocellulose membrane (Bio-Rad Inc. Hercules, CA, USA), blocked with Non-fat milk 5% in phosphate buffered saline (PBS)-Tween 0.05%, and incubated with anti-RBD rabbit polyclonal serum (1/1000 dilution). An anti-rabbit IgG horseradish-peroxidase conjugated (1/2000 dilution) was used as a secondary antibody (Agilent DAKO, Santa Clara, CA, USA) and 4-Chloro-1-Naphthol as substrate for colorimetric detection.

### ACE2 binding to RBD analyzed by titration ELISA

ACE2 binding to the produced RBD variant antigens was analyzed by ELISA. Briefly, 0.1ug of RBD per well was immobilized on a High binding Microplatte (Greiner Bio-One GmbH, Austria.) and incubated ON at 4°C. Next, plates were blocked with 5% non-fat milk in PBS 0.05% Tween20 for 2 h at 37°C and then washed with PBS and 0.05% Tween20. Plates were then incubated with hACE2 (R&D, Minneapolis MN, USA) for 2 h at 37°C and another washing step was performed. Finally, plates were incubated with anti-hACE2 (R&D Systems) diluted in 1% non-fat milk PBS 0.05% Tween 20 for 2 h at 37°C. Detection was performed with a secondary Ab conjugated to HRP (Agilent DAKO, Santa Clara, CA, USA) and visualized with tetramethylbenzidine (TMB) substrate (BD biosciences, Franklin Lakes, New Jersey, USA).

### Adjuvant and formulations

Pilot batches of vaccines were manufactured using aluminum hydroxide as adjuvant. The antigen was adsorbed onto the adjuvant by mixing. Free antigen was controlled to be less than 10% of total antigen in the vaccine.

### Ethics statement

All experimental protocols with animals were conducted in strict accordance with international ethical standards for animal experimentation (Helsinki Declaration and its amendments, Amsterdam Protocol of welfare and animal protection and National Institutes of Health, NIH USA, guidelines.). The protocols performed were also approved by the Institutional Committee for the use and care of experimental animals (CICUAE) from National University of San Martin (UNSAM) (01/2020).

### Animals and immunizations

Eight-week-old female BALB/c mice were obtained from IIB-UNSAM animal facility. Animals were intramuscularly (i.m) inoculated at day 0 and 14 with i) Gamma RBD (10μg) + Alum (n=5), ii) ancestral RBD (10μg) + Alum (n=5), iii) placebo (n=4). In other experiments, i) Gamma RBD (10μg) + Alum (n=5) and ii) placebo (n=5). Blood samples were collected weekly to measure total and neutralizing antibody titers and at day 42 post prime immunization animals were sacrificed and spleens and lungs harvested.

### Determination of antibody levels in serum

RBD-specific antibody responses (IgG) were evaluated by indirect ELISA as described previously^38^. Briefly, plates were coated with 0.1 μg/well of RBD (derived from ancestral or Gamma variant) in phosphate buffered saline (PBS) overnight at 4 °C. Plates were washed and then incubated with diluted sera and then plates were washed and incubated with HRP conjugated anti-mouse IgG (SIGMA, St. Louis, MO, USA). Results were read at 450 nm to collect end point ELISA data. End-point cut-off values for serum titer determination were calculated as the mean specific optical density (OD) plus 3 standard deviations (SD) from sera of saline immunized mice and titers were established as the reciprocal of the last dilution yielding an OD higher than the cut-off.

### Virusess

Ancestral SARS-CoV-2 (B.1, GISAID Accession ID EPI_ISL_16290469), Alpha (B.1.1.7 GISAID Accession ID EPI_ISL_15806335), Gamma (P.1, GISAID Accession ID EPI_ISL_15807444) and Omicron BA.5 (GISAID Accession ID EPI_ISL_16297058) were isolated from nasopharyngeal specimens at the Instituto de Investigaciones Biotecnológicas (IIB, UNSAM) and adapted in Vero E6 cultures. Delta SARS-CoV-2 (GISAID Accession ID: EPI_ISL_11014871) and Omicron BA.1 (GISAID Accession ID EPI_ISL_10633761) were isolated at Instituto de Investigaciones Biomédicas en Retrovirus y SIDA (INBIRS, UBA-CONICET) from nasopharyngeal swabs of patients. Studies using SARS-CoV-2 were done in a Biosafety level 3 laboratory at IIB, UNSAM, and the protocol was approved by the INBIRS Institutional Biosafety Committee.

### SARS-CoV-2 neutralization assay

Serum samples were heat-inactivated at 56°C for 30 min. Serial dilutions were performed and then incubated for 1 h at 37°C in the presence of 300 TCID_50_ of SARS-CoV-2 in DMEM 2% FBS. One hundred μl of the mixtures were then added onto Vero cells monolayers. After 72 h at 37°C and 5% CO_2_, cells were fixed with PFA 4% (4°C overnight) and stained with crystal violet solution in methanol. The cytopathic effect (CPE) of the virus on the cell monolayer was assessed by surface scanning at 585 nm in a microplate reader (FilterMAx F5 Microplate reader, Molecular Devices, San Jose, CA, USA). Average optical density (OD) of each well was used for the calculation of the percentage of neutralization of viral CPE for each sample as: Neutralization% = 100 x (DO _sample_ - DO _virus control_) / (DO _cell control_ - DO _virus control_). Non-linear curves of Neutralization (%) vs. Log 1/sera dilution were fitted to obtain the titer corresponding to the 50% of neutralization (NT50).

### Determination of T cell immune responses

Four weeks after the second dose, mice were sacrificed to study cellular responses. Splenocytes were cultured for 5 days in the presence of RBD antigen (ancestral or Gamma) or complete medium. Then, supernatants were collected, and cytokines (IFN-y and IL-5) were measured by ELISA (Biolegend. San Diego, CA). For intracellular cytokine determination: splenocytes were cultured (4×10^6^ cells/well) in the presence of stimulus medium (complete medium supplemented with anti-CD28 and anti-CD49d) or Ag stimuli (stimulus medium + RBD-peptides + RBD protein) for 18 h. Next, brefeldin A was added for 5 h to the samples. Cells were then washed, fixed, permeabilized, stained, and analyzed by flow cytometry. The cells were stained with Viability dye (Zombie Acqua), anti-mouse-CD8a Alexa Fluor 488, anti-mouse-CD4 Alexa Fluor 647, anti-IL-4 Brilliant Violet 421, anti-TNFα PeCy7 and anti-IFN-γ PE (Biolegend. San Diego, CA). BD Fortessa LSR-X Cytometer with DIVA Software were used for analysis.

### SARS-CoV-2 challenge of vaccinated K18-hACE2 mice

Four-week-old K18-hACE2 mice from Jackson Laboratory were used for evaluating vaccine efficacy. Mice were separated into two groups: i) control (n=8) Placebo (PBS) and Gamma RBD +Alum (n=9). Mice in each group included males and females. They were i.m immunized on day 0 and 14 as described for immunogenicity assays. Four weeks post second vaccination, mice were challenged intranasally (i.n.) with 10^5^ PFU of SARS-CoV-2 strain WA1/2010 split evenly in each nare. Mice were monitored daily for weight loss and signs of disease for two weeks post-challenge. Three mice per group were euthanized on day 5 post-challenge to evaluate organ viral loads, by plaque assay on Vero E6 cells as previously described^38^.

### Statistical analysis

Statistical analysis and plotting were performed using GraphPad Prism 8 software (GraphPad Software, San Diego, CA). In experiments with more than two groups, data were analyzed using one-way ANOVA with Kruskal Wallis or Bonferroni post-test. When necessary, a logarithmic transformation was applied prior to the analysis to obtain data with a normal distribution. In experiments with two groups, an unpaired t test or Mann–Whitney U test were used. A p value <0.05 was considered significant. When bars were plotted, results were expressed as means ± SEM for each group. Survival curves were analyzed using Long-rank test.

## Data availability

The authors declare that the data supporting the findings of this study are available from the corresponding author upon reasonable request.

## Acknowledgements

The authors are grateful to the technical staff at the I+D Biofármacos Laboratorio Pablo Cassará and to Fundación Pablo Cassará for their significant assistance in the preparation and control of the antigen pilot lots and the regulatory documentation: Cintia Parsza, Brenda Heinrich, Melisa Gambone, Horacio Descoins, Christian Cortez, María Victoria Román and Romina Albarracín. The authors would also like to thank Danielle Porier, Krisangel López, and Manette Tanelus for technical assistance with the SARS-CoV-2 challenge studies. We thank Ministerio de Salud de la Provincia de Buenos Aires for providing mRNA, adenoviral and inactivated vaccines.

## Contributions

L.M.C. was responsible of overall experiment design, conducted humoral and cellular experiments, collected data, performed data analysis, and wrote the manuscript. J.C. was responsible for overall experimental design and conceptualization, wrote the manuscript and received funding support. J.M.R. generated high expression vectors, designed the antigen and was responsible of development of the vaccine formulation and downstream development. Coordination of antigen and vaccine production. K.A.P. and J.M.F. participated of experimental design, data analysis and reviewed the manuscript. D.E.A. designed and supervised neutralization studies and analysis. A.D. conducted and analyzed humoral studies and conducted cellular studies. C.P.C., L.A.B, L. M.S and L.P. conducted humoral and cellular studies. M.R.M. and E.F.C. performed virus neutralization studies and data analysis. S.A.DP. generated high expression vector and optimization of high productive clones. A.C.HI. Responsible of upstream and downstream development of antigens. I.G.K. coordinated overall analytics procedures, developed validation and analytics methodologies, performed structural characterization of antigen and vaccine formulation. A.V.DN. generated cell banks, performed stability, clones purity and identity studies. V.K. and I.D. generated, characterized, and screened cell clones. F.M.Z., J.A.B. and C.J.P performed upstream and downstream development of antigen. J.E.S. and M.LC. participated in the development of validation and analytics methodologies, performed structural characterization of antigen and vaccine formulation and performed animals studies. J.C.V. participated in the development of vaccine formulation, collaborates in the overall process of vaccine development. A.J.A. design, supervise and conduct animal challenge studies and data analysis. W.B.S. performed animal challenge studies and data analysis. All authors revised the manuscript.

## References

1 Baden, L. R. et al. Efficacy and Safety of the mRNA-1273 SARS-CoV-2 Vaccine. New England Journal of Medicine 384, 403–416, doi:10.1056/nejmoa2035389 (2021).

2 Bos, R. et al. Ad26 vector-based COVID-19 vaccine encoding a prefusion-stabilized SARS-CoV-2 Spike immunogen induces potent humoral and cellular immune responses. npj Vaccines 5, doi:10.1038/s41541-020-00243-x (2020).

3 Polack, F. P. et al. Safety and Efficacy of the BNT162b2 mRNA Covid-19 Vaccine. New England Journal of Medicine 383, 2603–2615, doi:10.1056/nejmoa2034577 (2020).

4 Voysey, M. et al. Safety and efficacy of the ChAdOx1 nCoV-19 vaccine (AZD1222) against SARS-CoV-2: an interim analysis of four randomised controlled trials in Brazil, South Africa, and the UK. The Lancet 397, 99–111, doi:10.1016/s0140-6736(20)32661-1 (2021).

5 Al Kaabi, N. et al. Effect of 2 Inactivated SARS-CoV-2 Vaccines on Symptomatic COVID-19 Infection in Adults. JAMA 326, 35, doi:10.1001/jama.2021.8565 (2021).

6 Chemaitelly, H. et al. Waning of BNT162b2 Vaccine Protection against SARS-CoV-2 Infection in Qatar. New England Journal of Medicine 385, e83, doi:10.1056/nejmoa2114114 (2021).

7 Goldberg, Y. et al. Waning Immunity after the BNT162b2 Vaccine in Israel. New England Journal of Medicine 385, e85, doi:10.1056/nejmoa2114228 (2021).

8 Wang, Q. et al. Antibody evasion by SARS-CoV-2 Omicron subvariants BA.2.12.1, BA.4 and BA.5. Nature 608, 603–608, doi:10.1038/s41586-022-05053-w (2022).

9 Khan, K. et al. Omicron BA.4/BA.5 escape neutralizing immunity elicited by BA.1 infection. Nature Communications 13, doi:10.1038/s41467-022-32396-9 (2022).

10 Tuekprakhon, A. et al. Antibody escape of SARS-CoV-2 Omicron BA.4 and BA.5 from vaccine and BA.1 serum. Cell 185, 2422–2433.e2413, doi:10.1016/j.cell.2022.06.005 (2022).

11 Tseng, H. F. et al. Effectiveness of mRNA-1273 against SARS-CoV-2 Omicron and Delta variants. Nature Medicine 28, 1063–1071, doi:10.1038/s41591-022-01753-y (2022).

12 Suryawanshi, R. K. et al. Limited cross-variant immunity from SARS-CoV-2 Omicron without vaccination. Nature 607, 351–355, doi:10.1038/s41586-022-04865-0 (2022).

13 Chalkias, S. et al. A Bivalent Omicron-Containing Booster Vaccine against Covid-19. New England Journal of Medicine 387, 1279–1291, doi:10.1056/nejmoa2208343 (2022).

14 Akache, B. et al. Immunogenicity of SARS-CoV-2 spike antigens derived from Beta & amp; Delta variants of concern. npj Vaccines 7, doi:10.1038/s41541-022-00540-7 (2022).

15 Sridhar, S. et al. The potential of Beta variant containing COVID booster vaccines for chasing Omicron in 2022. Nature Communications 13, doi:10.1038/s41467-022-33549-6 (2022).

16 Launay, O. et al. Immunogenicity and Safety of Beta-Adjuvanted Recombinant Booster Vaccine. New England Journal of Medicine 387, 374–376, doi:10.1056/nejmc2206711 (2022).

17 Sir Karakus, G. et al. Preclinical efficacy and safety analysis of gamma-irradiated inactivated SARS-CoV-2 vaccine candidates (Cold Spring Harbor Laboratory, 2020).

18 Turan, R. D. et al. Gamma-irradiated SARS-CoV-2 vaccine candidate, OZG-38.61.3, confers protection from SARS-CoV-2 challenge in human ACEII-transgenic mice. Scientific Reports 11, doi:10.1038/s41598-021-95086-4 (2021).

19 Khorattanakulchai, N. et al. Receptor binding domain proteins of SARS-CoV-2 variants produced in *Nicotiana benthamiana* elicit neutralizing antibodies against variants of concern. Journal of Medical Virology, doi:10.1002/jmv.27881 (2022).

20 Winkler, E. S. et al. SARS-CoV-2 infection of human ACE2-transgenic mice causes severe lung inflammation and impaired function. Nature Immunology 21, 1327–1335, doi:10.1038/s41590-020-0778-2 (2020).

21 Jiang, R.-D. et al. Pathogenesis of SARS-CoV-2 in Transgenic Mice Expressing Human Angiotensin-Converting Enzyme 2. Cell 182, 50–58.e58, doi:10.1016/j.cell.2020.05.027 (2020).

22 Harvey, W. T. et al. SARS-CoV-2 variants, spike mutations and immune escape. Nat Rev Microbiol 19, 409–424, doi:10.1038/s41579-021-00573-0 (2021).

23 Tao, K. et al. The biological and clinical significance of emerging SARS-CoV-2 variants. Nat Rev Genet 22, 757–773, doi:10.1038/s41576-021-00408-x (2021).

24 Zhao, X. et al. Neutralisation of ZF2001-elicited antisera to SARS-CoV-2 variants. Lancet Microbe 2, e494, doi:10.1016/S2666-5247(21)00217-2 (2021).

25 Brinkkemper, M. et al. A third SARS-CoV-2 spike vaccination improves neutralization of variants-of-concern. NPJ Vaccines 6, 146, doi:10.1038/s41541-021-00411-7 (2021).

26 Branche, A. R. et al. SARS-CoV-2 Variant Vaccine Boosters Trial: Preliminary Analyses (Cold Spring Harbor Laboratory, 2022).

27 Wang, Z. et al. A potent, broadly protective vaccine against SARS-CoV-2 variants of concern. NPJ Vaccines 7, 144, doi:10.1038/s41541-022-00571-0 (2022).

28 Chalkias, S. et al. Safety, immunogenicity and antibody persistence of a bivalent Beta-containing booster vaccine against COVID-19: a phase 2/3 trial. Nat Med 28, 2388–2397, doi:10.1038/s41591-022-02031-7 (2022).

29 Dai, L. et al. Efficacy and Safety of the RBD-Dimer-Based Covid-19 Vaccine ZF2001 in Adults. N Engl J Med 386, 2097–2111, doi:10.1056/NEJMoa2202261 (2022).

30 Xu, K. et al. Protective prototype-Beta and Delta-Omicron chimeric RBD-dimer vaccines against SARS-CoV-2. Cell 185, 2265–2278 e2214, doi:10.1016/j.cell.2022.04.029 (2022).

31 Grigoryan, L. et al. Adjuvanting a subunit SARS-CoV-2 vaccine with clinically relevant adjuvants induces durable protection in mice. NPJ Vaccines 7, 55, doi:10.1038/s41541-022-00472-2 (2022).

32 Gilbert, P. B. et al. A Covid-19 Milestone Attained - A Correlate of Protection for Vaccines. N Engl J Med, doi:10.1056/NEJMp2211314 (2022).

33 Moss, P. The T cell immune response against SARS-CoV-2. Nature Immunology 23, 186–193, doi:10.1038/s41590-021-01122-w (2022).

34 Guerrera, G. et al. BNT162b2 vaccination induces durable SARS-CoV-2-specific T cells with a stem cell memory phenotype. Sci Immunol 6, eabl5344, doi:10.1126/sciimmunol.abl5344 (2021).

35 Swanson, P. A., 2nd et al. AZD1222/ChAdOx1 nCoV-19 vaccination induces a polyfunctional spike protein-specific T(H)1 response with a diverse TCR repertoire. Sci Transl Med 13, eabj7211, doi:10.1126/scitranslmed.abj7211 (2021).

36 Jara, A. et al. Effectiveness of homologous and heterologous booster doses for an inactivated SARS-CoV-2 vaccine: a large-scale prospective cohort study. Lancet Glob Health 10, e798–e806, doi:10.1016/S2214-109X(22)00112-7 (2022).

37 Rouco, S. O. et al. Heterologous booster response after inactivated virus BBIBP-CorV vaccination in older people. Lancet Infect Dis 22, 1118–1119, doi:10.1016/S1473-3099(22)00427-3 (2022).

38 Coria, L. M. et al. A Novel Bacterial Protease Inhibitor Adjuvant in RBD-Based COVID-19 Vaccine Formulations Containing Alum Increases Neutralizing Antibodies, Specific Germinal Center B Cells and Confers Protection Against SARS-CoV-2 Infection in Mice. Front Immunol 13, 844837, doi:10.3389/fimmu.2022.844837 (2022).

